# Morphology of the nervous system of monogonont rotifer *Epiphanes senta* with focus on sexual dimorphism between feeding females and dwarfed males

**DOI:** 10.1101/643817

**Authors:** Ludwik Gąsiorowski, Anlaug Furu, Andreas Hejnol

## Abstract

**Background:** Monogononta is a large clade of rotifers comprised of diverse morphological forms found in a wide range of ecological habitats. Most of the monogonont species display a cyclical parthenogenesis, where generations of asexually reproducing females are interspaced by mixis events when sexual reproduction occurs between mictic females and dwarfed, haploid males. The morphology of monogonont feeding females is relatively well described, however data on male anatomy are very limited. Thus far, male musculature of only two species has been described with confocal laser scanning microscopy (CLSM) and it remained unknown how dwarfism influences neuroanatomy of males.

**Results:** Here, we provide a CLSM-based description of the nervous system of both sexes of *Epiphanes senta*, a freshwater monogonont rotifer. The general nervous system architecture is similar between males and females and shows same level of complexity. However, the nervous system in males is more compact and lacks its stomatogastric part.

**Conclusion:** Comparison of the neuroanatomy between male and normal-sized feeding females provides better understanding of the nature of male dwarfism in Monogononta. We propose that dwarfism of monogonont non-feeding males is a specific case of progenesis as they, due to their inability to feed, retain a juvenile body size. Reduction of the stomatogastric nervous system in the males correlates with the loss of entire digestive tract and associated morphological structures.

## Background

Monogononta is a large clade belonging to Rotifera (=Syndermata) with about 1600 species formally described(1). These microscopic animals inhabit both freshwater and marine environments, and occupy many different ecological niches from being sessile suspension feeders to active planktonic predators(1, 2). This ecological diversity is coupled with a vast variety of body plans(3) and morphological adaptations to their particular life style. Despite this variation of monogonont morphology, it is often possible to distinguish three main body regions: 1. head, equipped with a wheel organ or corona, which serves for food capture and locomotion, 2. trunk, which contains, among other organs, the characteristic pharynx (mastax) with sclerotized jaws (trophi) and 3. posterior foot with terminal paired toes where pedal glands that are used for adhesion to the substrate are localized(1). Similarly to bdelloids, another large rotiferan clade, monogononts are able to reproduce asexually by lying parthenogenetic eggs. Under normal conditions this type of reproduction dominates(2, 4, 5). However, unlike bdelloids which are exclusively parthenogenetic, most monogonont species also reproduce sexually, often as a response to stressful environmental conditions(2, 4-9). The monogonont haploid males are predominantly dwarfed and short-living, often with a reduced digestive system with a single testicle and copulatory organs occupying most of their body(5, 10-12).

Nervous system architecture has been studied in many monogonont species from diverse evolutionary lineages and ecological niches using light microscopy and TEM as well as histochemical and immunohistochemical techniques combined with epifluorescent and confocal laser scanning microscopy (e.g.(12-33)). Further, gene expression in the developing and juvenile nervous system of monogonont rotifers has recently been studied(34, 35). However, most of these studies focused on the nervous system of feeding females, whereas neuroanatomy of dwarfed males remains poorly examined. The only available information on the male nervous system dates back to the light microscopy investigation of Remane(31) and a single histofluorescent labeling of the catecholaminergic structures combined with epifluorescent light microscopy(12). Neither of these studies provide great resolution of examined structures or detailed comparison of male and female neuroanatomies. So far, the only work that systematically treated sexual dimorphism in monogonont morphology focused on body musculature(11). Therefore, it remains unknown how male dwarfism influences nervous system architecture in Monogononta.

*Epiphanes senta* (Müller, 1773) was one of the species investigated for sexual dimorphism in musculature by Leasi et al.(11). It is a relatively large freshwater rotifer, found around the world in littoral habitats of eutrophic water bodies, such as lakes, small ponds, astatic pools and floodplains(9, 36, 37). Females are relatively stationary, mostly attached or slowly swimming near the substrate, feeding on algae and bacteria, which they filtrate using the corona ((11, 37), personal observation). However, they can also ascend to the water column and cases of cannibalism have been observed ((11), personal observation). The males are much smaller than females(11, 36). They can be found at all times, although normally in small densities in the animal lab cultures (personal observation). Further, the males of *E. senta* display a unique precopulatory mating behavior – the male seems to sense and prioritizes eggs of prospective mictic females and then copulates with the female as she emerges from the egg(9, 37).

In order to test if male dwarfism is coupled with substantial changes in the neuroanatomy we investigated the nervous system of females and dwarf males of *E. senta* using confocal laser scanning microscopy (CLSM) combined with the antibody staining against common nervous system markers (tyrosinated tubulin, acetylated tubulin, serotonin and FMRF-amide). Accordingly, we provide a CLSM-based detailed description of nervous system of monogonont dwarfed males. By comparing it to the nervous system of conspecific females we can better understand the nature of male dwarfism in Monogononta, as well as infer impact of this phenomenon on the morphology of one of the most crucial organ systems.

## Results

### Taxonomical remark

Schröder and Walsh(36) reported that *E. senta* is a species complex of morphologically almost identical cryptic species which mostly differ from each other in geographical distribution, details of trophi morphology and the sculpturing of the resting egg shell. We assume that animals, which we used in our study, represent *E. senta*, however for the sake of future exact taxonomical identification we searched the transcriptome of the investigated species for COX1 sequence. We obtained two sequences of 686 bp each, which differ between each other in 6 nucleotides (either due to intraspecific polymorphism or sequencing inaccuracy) and are available as Additional file 1.

### General morphology

The body of both sexes of *E. senta* is clearly divided into three regions: head with corona, trunk and foot (Fig. 1). Males and fully developed females clearly differ in body size (Fig. 1A, C), with the mean body length of ≈220μm (N=3) for males and ≈487μm (N=7) for females. However, newly hatched females are substantially smaller (the smallest measured specimen was 340μm long) and could be confused with males if solely considered by body size. Due to the fact that the body wall of *E. senta* is transparent it is possible to identify most of the internal organs, including gonads, glands and protonephridial terminal organs, using light microscopy (Fig. 1A, B). Even though body shape and proportions are similar between the sexes, males obviously lack any elements of the digestive tract (Fig. 1B). Additionally, a single testicle with individual spermatozoa visible in LM (*te*, Fig. 1B) is found in the posterior part of the male trunk which makes it easy to distinguish between the sexes regardless of the body size.

**Fig. 1.**
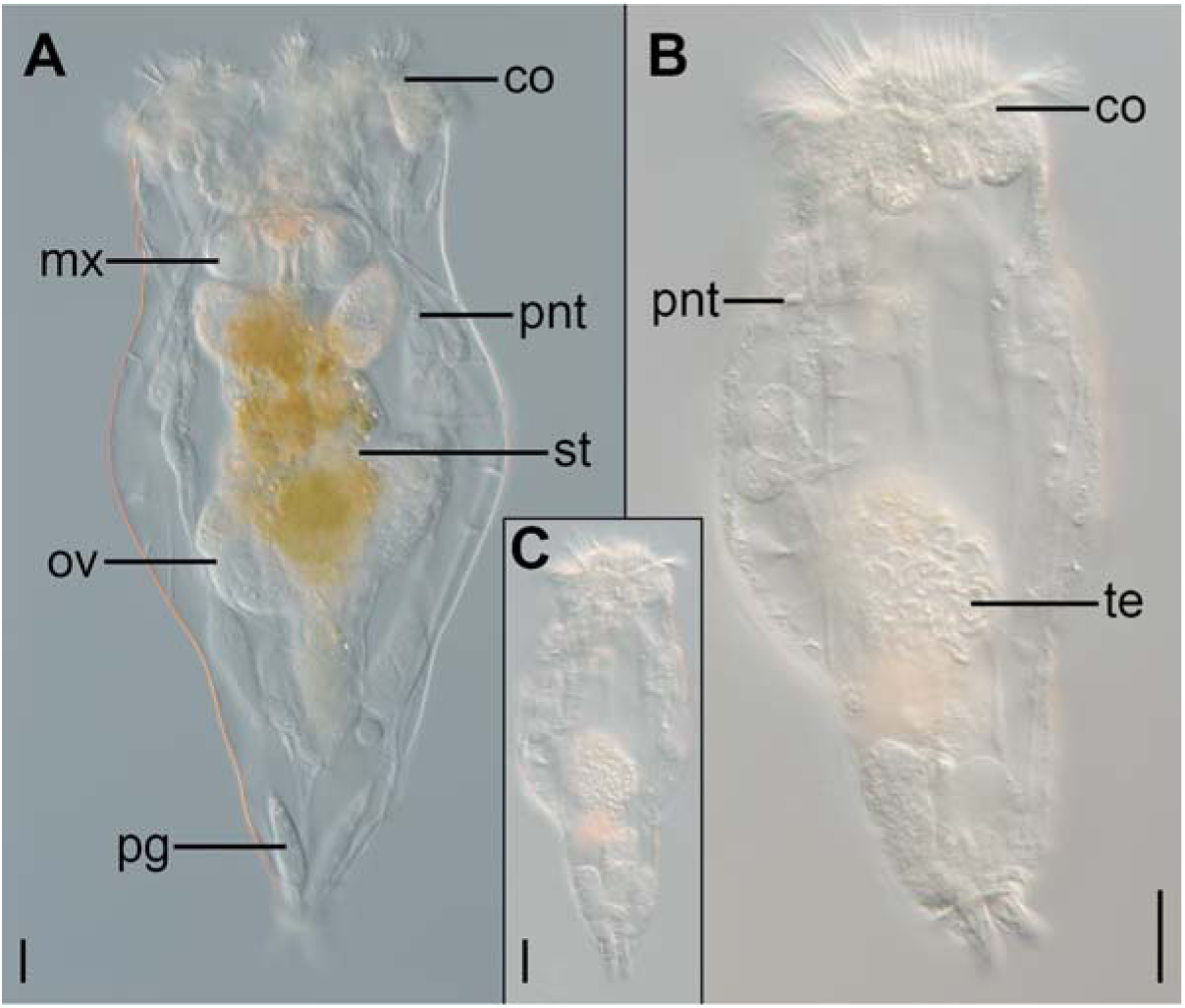
Light micrographs showing sexual dimorphism in *Epiphanes senta*. **A** female, **B** enlarge picture of the male, **C** male in scale to the female. Scale bar on all panels 20 μm. Abbreviations: *co* corona, *mx* mastax, *ov* ovary, *pg* pedal gland, *pnt* protonephridial terminal organ, *st* stomach, *te* testes with spermatozoa.

### Nervous system of the female

The nervous system of feeding females consists of 1) the brain, located in the dorso-posterior part of the head, 2) two longitudinal nerve cords extending laterally along the trunk, connected by two commissures and merging posteriorly in the foot ganglion, 3) coronal nerves, 4) peripheral nerves and sensory organs, and 5) stomatogastric nervous system related with mastax (Figs. 2A–C; 3A; 4A–C; 5A, B).

**Fig. 2.**
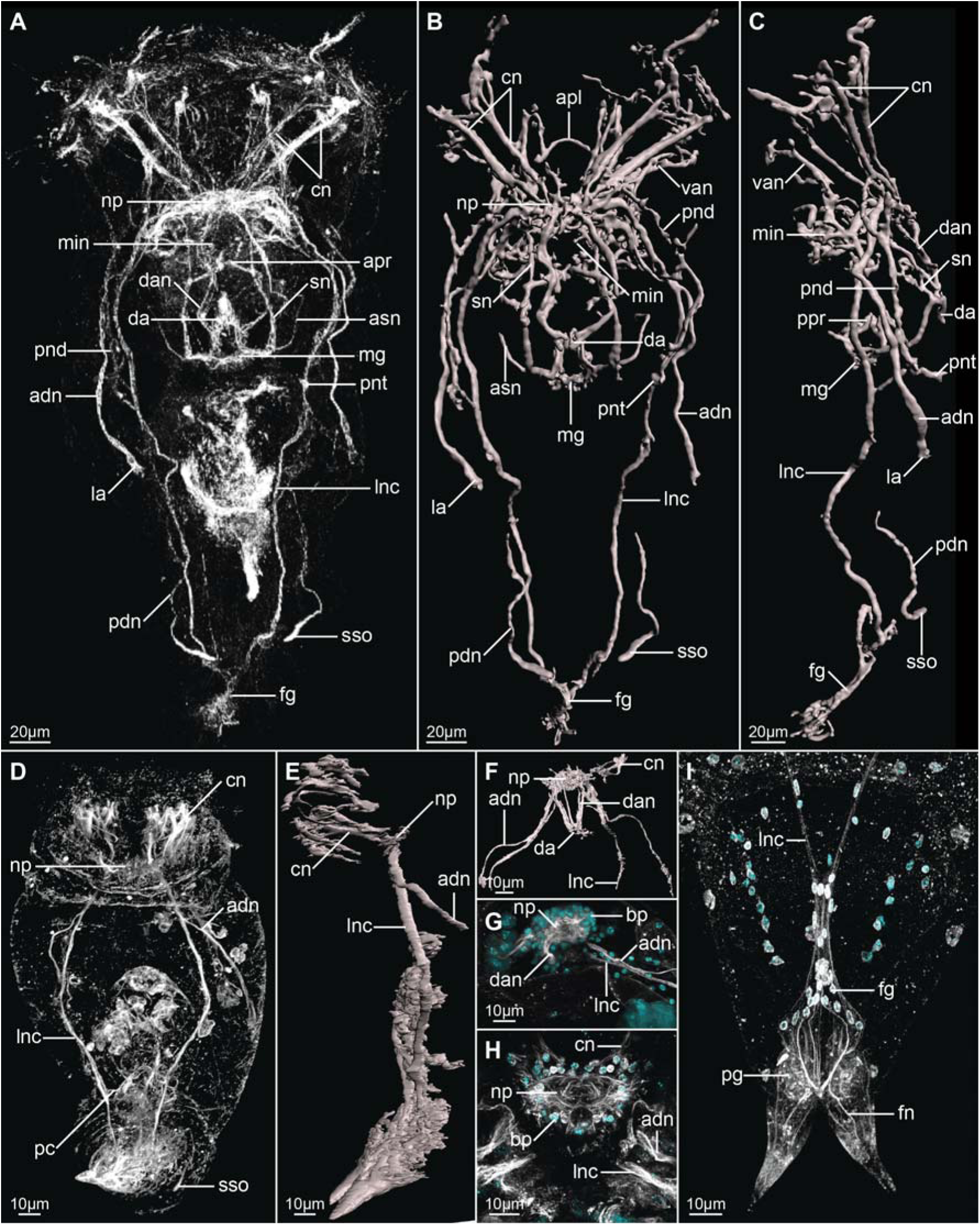
Z-projections (**A, D, G**–**I**) and 3-D reconstructions (**B, C, E** and **F**) of the nervous system of *Epiphanes senta* females (**A**–**C, H, I**) and males (**D**–**G**), visualized with CLSM combined with antibody staining against tyrosinated-tubulin (white) and DAPI staining of cell nuclei (cyan). Entire animals in dorso-ventral (**A, B, D**) and lateral (**C, E**) views. Details of the anterior part of the nervous system (**F**), brain (**G, H**) and posterior structures (**I**). In all panels anterior is to the top. Abbreviations: *adn* anterior dorsal nerve, *apl* anterior protonephridial loop, *apr* anterior pharyngeal receptor, *asn* accessory stomatogastric nerve, *bp* brain perikarya, *cn* coronal nerves, *da* dorsal antenna, *dan* nerve of dorsal antenna, *fg* foot ganglion, *fn* foot nerve, *la* lateral antenna, *lnc* longitudinal nerve cord, *mg* mastax ganglion, *min* mouth innervation, *np* neuropile, *pc* posterior commissure, *pdn* posterior dorsal nerve, *pg* pedal gland, *pnd* protonephridial duct, *pnt* protonephridial terminal organ, *ppr* posterior pharyngeal receptor, *sn* stomatogastric nerve, *sso* supraanal sensory organ, *van* ventro-anterior nerve.

**Fig. 3.**
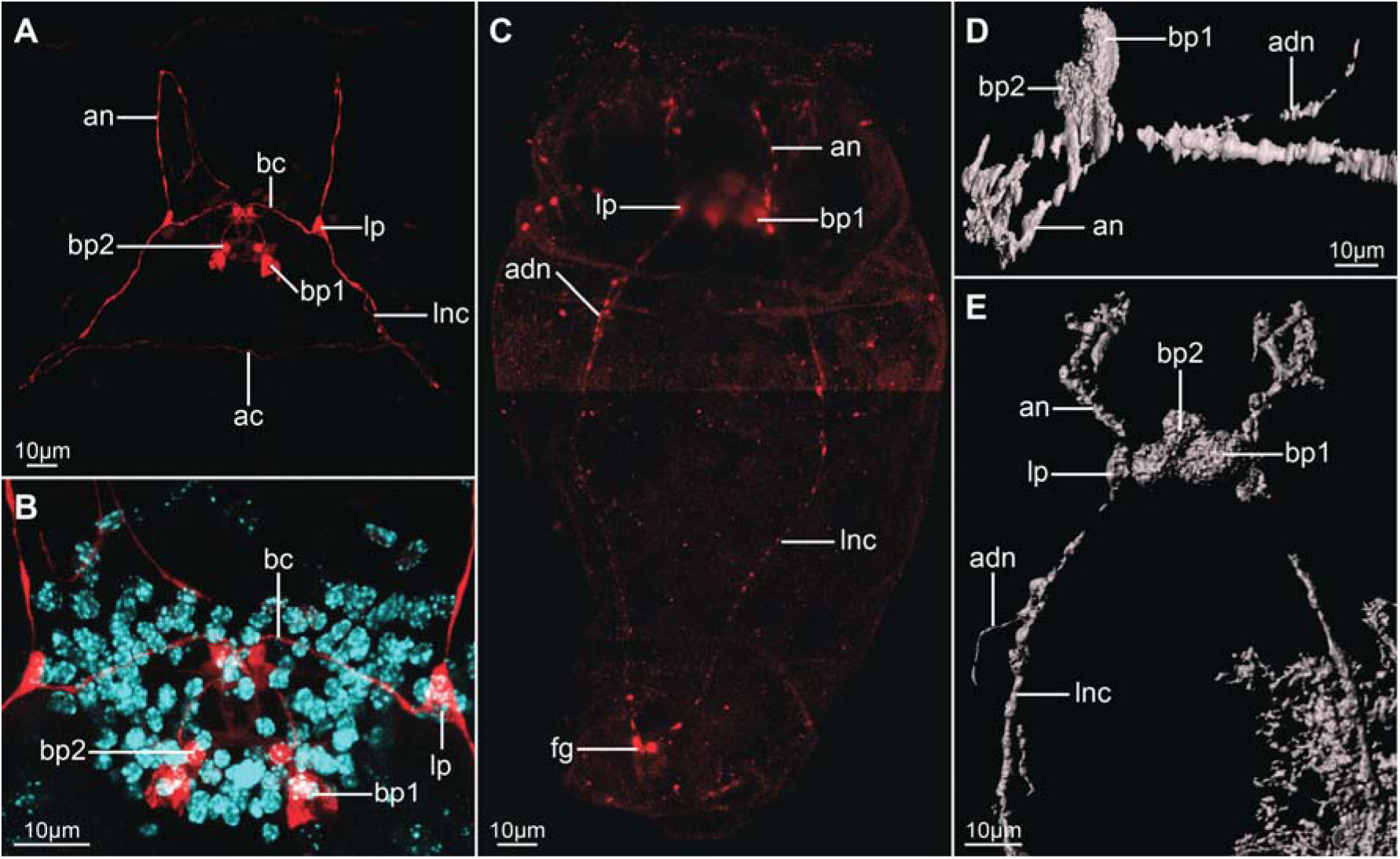
Serotonin-like immunoreactivity in the nervous system of *Epiphanes senta* females (**A, B**) and males (**C**–**E**). Z-projections of CLSM (**A**–**C**) showing antibody staining against serotonin (red) and DAPI staining of cell nuclei (cyan) and 3-D reconstructions (**D, E**) in dorso-ventral (**A**–**C, E**) and lateral (**D**) views. Details of the anterior part of the nervous system (**A, D, E**) and brain (**B**). Anterior is to the top (**A**–**C, E**) and to the left (**D**), dorsal to the top on panel **D**. Abbreviations: *ac* anterior commissure, *adn* anterior dorsal nerve, *an* anterior nerve, *bc* brain commissure, *bp* brain perikarya, *fg* foot ganglion, *lnc* longitudinal nerve cord, *lp* lateral perikaryon.

**Fig. 4.**
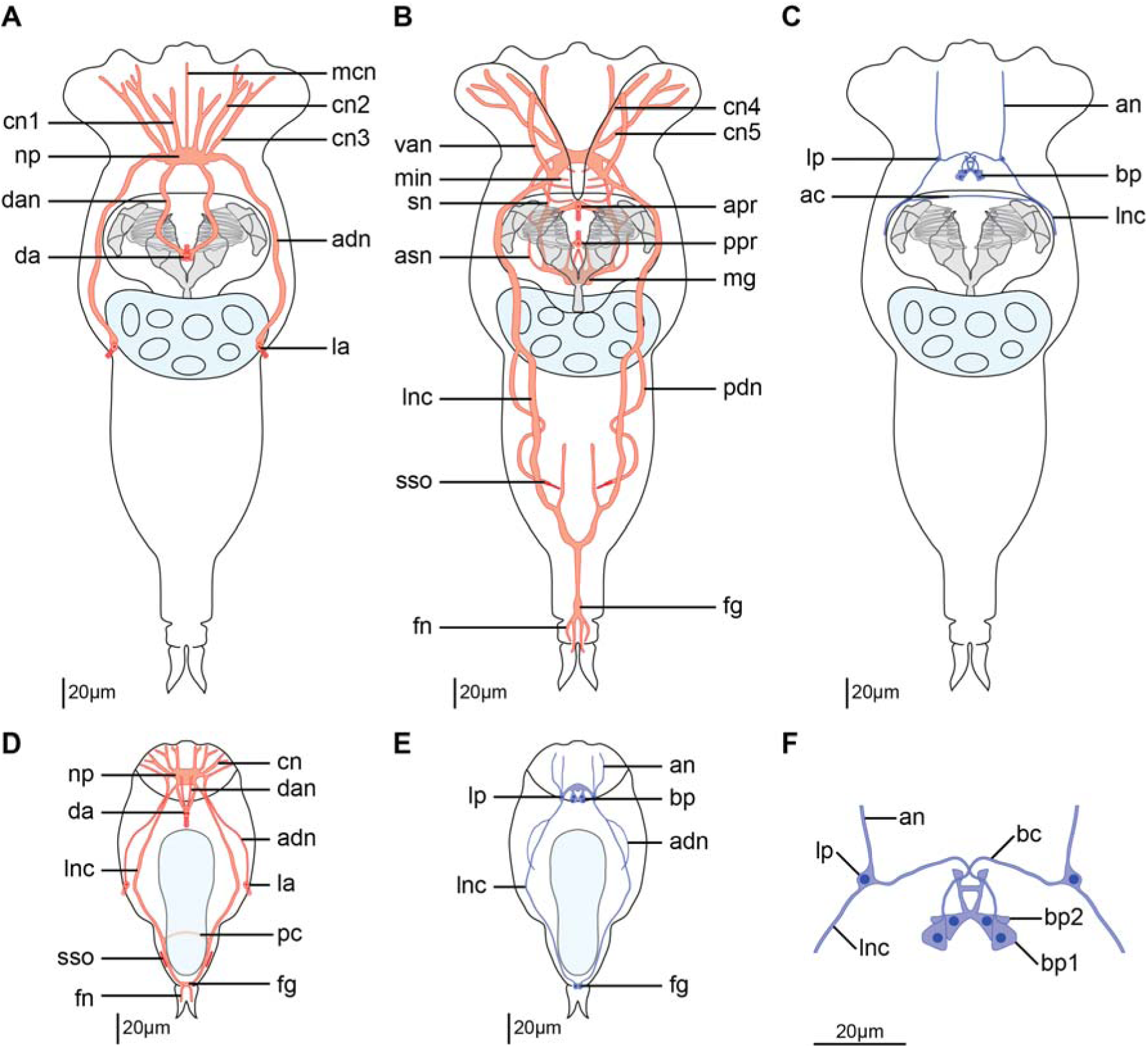
Schematic drawings of the nervous system of *Epiphanes senta* females (**A**–**C, F**) and males (**D, E**) inferred from tyrosinated tubulin-like immunoreactivity (red) and serotonin-like immunoreactivity (dark blue). Dorsal structures (**A**), ventral structures (**B**), entire body (**C**–**E**) and details of the brain (**F**) in dorso-ventral view with anterior to the top. Abbreviations: *ac* anterior commissure, *an* anterior nerve, *and* anterior dorsal nerve, *apr* anterior pharyngeal receptor, *asn* accessory stomatogastric nerve, *bc* brain commissure, *bp* brain perikarya, *cn* coronal nerves, *dan* nerve of dorsal antenna, *fg* foot ganglion, *fn* foot nerve, *la* lateral antenna, *lnc* longitudinal nerve cord, *lp* lateral perikaryon, *mcn* median coronal nerve, *mg* mastax ganglion, *min* mouth innervation, *np* neuropile, *pc* posterior commissure, *pdn* posterior dorsal nerve, *ppr* posterior pharyngeal receptor, *sn* stomatogastric nerve, *sso* supraanal sensory organ, *van* ventro-anterior nerve.

**Fig. 5.**
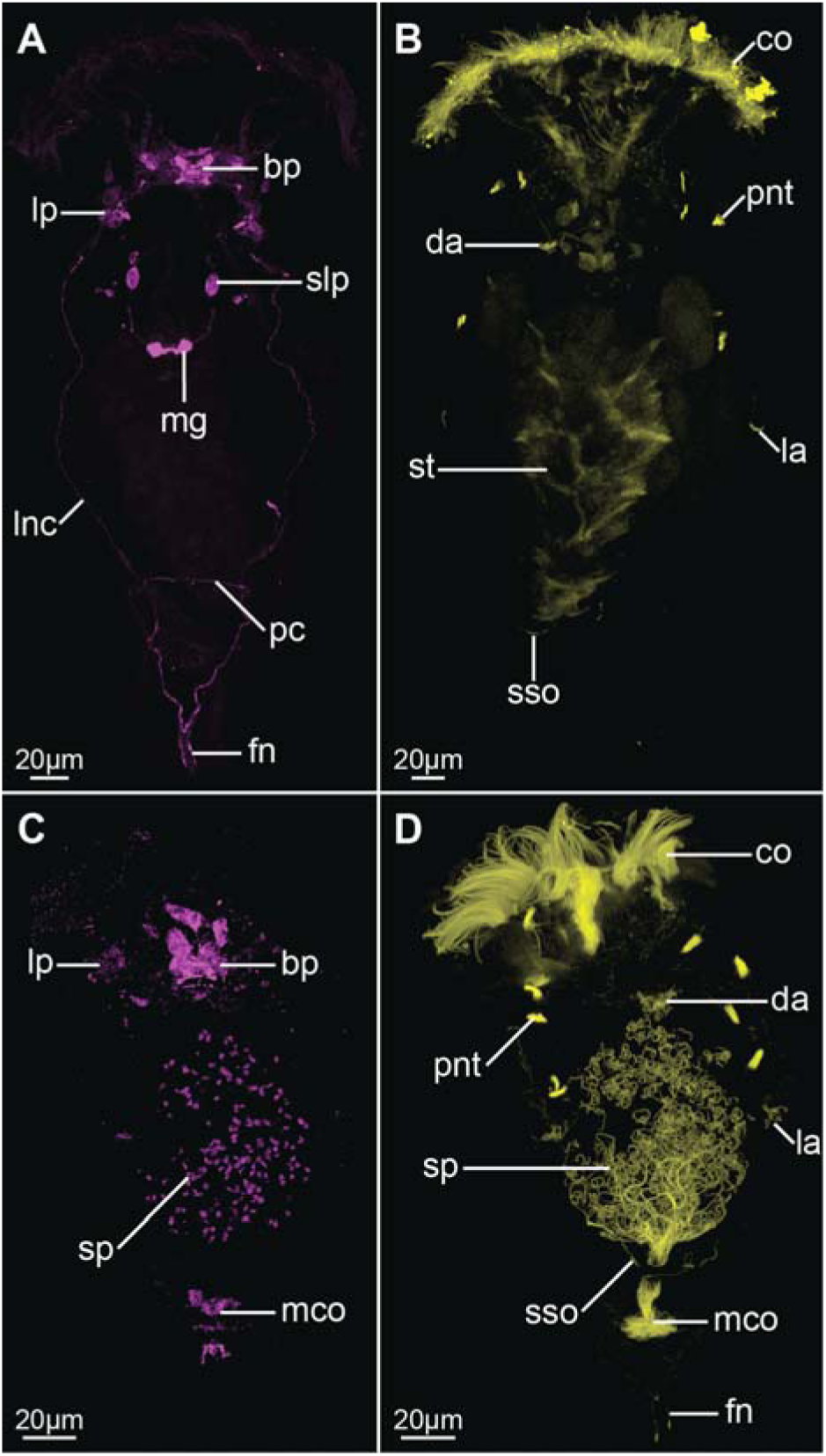
Z-projections showing FMRF-amide-like (**A, C**) and acetylated-tubulin-like (**B, D**) immunoreactivity in *Epiphanes senta* females (**A, B**) and males (**C, D**). Dorso-ventral view with anterior to the top on all panels. Abbreviations: bp brain perikarya, *co* corona, *da* dorsal antenna, *fn* foot nerve, *la* lateral antenna, *lnc* longitudinal nerve cord, *lp* lateral perikarya, *mco* male copulatory organs, *mg* mastax ganglion, *pc* posterior commissure, *pnt* protonephridial terminal organ, *slp* stomatogastric lateral perikaryon, *sp* spermatozoa, *sso* supraanal sensory organ, *st* stomach.

The brain is ellipsoidal (mean length ≈24μm, mean width ≈58μm; N=7) and consists of an external layer of perikarya and internal neuropile (*bp* and np, Fig. 2H, respectively). Anteriorly and directly from the brain 11 coronal nerves (5 paired and one single median nerve) originate (*cn* and *mcn*, Figs. 2A–C, H; 4 A, B) and innervates large, cushion-shaped cells at the edge of corona (both in trochus and cingulum). Laterally, two thick bundles of neurites emerge from the brain; one of them (*adn*) is more dorsal and leads to the lateral antennae (*la*). The second bundle gives rise to the thick longitudinal nerve cords (*lnc*), fine neurites innervating mouth opening (*min*) and ventral anterior nerves (*van*), which extend to the ventral part of the corona (Figs. 2A–C, H; 3A, B). Two pairs of nerves connect to the posterior brain: ventrally the stomatogastric nerves (*sn*), and dorsally the nerves of the dorsal antenna (*dan*) (Figs. 2A–C; 3A, B). Staining against serotonin revealed presence of three pairs of serotonin-like immunoreactive (SLIR) perikarya in the brain of female (Fig. 3A, B; 4F), two of which form clusters in the dorso-posterior part of the brain (*bp1* and 2, Figs. 3A, B; 4F). Neurites of BP1 extends contralaterally, cross each other in the anterior brain, forming the only SLIR commissure of the brain (*bc*) and then connect the lateral SLIR perikarya (*lp*, Figs. 3A, B; 4F). From this point one SLIR neurite (*an*) extends anteriorly to the corona (note it is not identical with any of the aforementioned coronal nerves) and the second SLIR neurite (*lnc*) contribute to the lateral nerve cord (Figs. 3A; 4F). FMRF-amide-like immunoreactivity (FLIR) was detected as well in some of the brain perikarya (*bp*, Fig. 5A). However, it was impossible to determine the exact number and connectivity of FLIR neurons.

Lateral nerve cords (*lnc*, Figs. 2A–C; 3A; 4B, C, 5A) extend along the trunk and posteriorly they merge in the foot ganglion (*fg*), which is a concentration of around 25 perikarya located at the trunk/foot boundary (Fig. 2I). Short nerves extend from the foot ganglion towards pedal glands and tips of the foot toes (*fn*, Figs. 2I, 4B, 5A). At the level of the gonad a single posterior dorsal nerve (*pdn*) originates from each nerve cord and extends dorsally (Figs. 2A–C, 4B). This meandering bundle eventually innervates the supra-anal sensory organ (*sso*) in the dorso-posterior part of the trunk (Figs. 2A–C; 4B; 5B). Some of the neurites of the lateral nerve cords are SLIR (only in the anterior portion of the cord, Figs. 3A, 4C) and FLIR (Figs. 5A). The SLIR neurites form anterior commissure connecting longitudinal cords ventrally at the level of the anterior mastax, whereas FLIR neurites form a posterior commissure at the level of hindgut (*ac*, Figs. 3A; 4C and *pc*, Fig. 5A, respectively). There are clusters of several FLIR perikarya related with the anterior section of each nerve cord laterally to the mouth opening (*lp*, Fig. 5A), but there are no SLIR perikarya related with the lateral nerve cords.

The stomatogastric nervous system (SNS) makes a large portion of the female nervous system. The mastax ganglion (*mg*) is a central element of the SNS located in the posterior part of the mastax (Figs. 2A–C, 4B; 5A). Tyrosinated tubulin-like immunoreactivity was detected in the central portion of the ganglion, whereas two of its perikarya are FLIR. A pair of stomatogastric nerves (*sn*) connects mastax ganglion with the ventro-posterior brain and give rise to short accessory stomatogastric nerves (*asn*) innervating lateral portion of the mastax (Figs. 2A–C, 4B). At least one pair of large FLIR perikarya is present along stomatogastric nerves (*slp*, Fig. 5A). Two pharyngeal unicellular ciliated receptors are associated with SNS: one (*apr*) in the anterior mastax, with cilia protruding posteriorly, and the second (*ppr*) in the posterior mastax with cilia protruding anteriorly, between jaws (Figs. 2A–C; 4B). The anterior pharyngeal receptor connects to the stomatogastric nerves, whereas posterior one is innervated directly from the mastax ganglion. There are no SLIR structures in the SNS of *E. senta* females.

Staining against tyrosinated and acetylated tubulin revealed five sensory organs on the surface of the female body that connect with the nervous system. Three of them (unpaired dorsal antenna in the posterior head and paired lateral antennae in the middle portion of the trunk) are multiciliated cells that protrude their cilia to the external environment (*da* and *la*, Figs. 2A–C; 4A; 5B). There is at least one cell nucleus related with each lateral antenna and two cell nuclei related with the dorsal antenna. The second type of sensory organs is represented by paired supra-anal sensory organs positioned laterally to the anal opening on the dorsal side of the body (*sso*, Figs. 2A–C; 4B; 5B). The individual cilia are never visible in the organ and it appears instead as a solid, elongated structure with a strong immunoreactivity, which seems to be directly continuous with the posterior dorsal nerve. There is also no evident nucleus related with the supra-anal sensory organ.

Further, part of the excretory system was stained with the antibodies against tyrosinated and acetylated tubulin. Acetylated tubulin-like immunoreactivity was detected in the four pairs of ciliated terminal organs of the protonephridial system (*pnt*, Fig. 5B), showing a typical monogonont organization with all particular cilia of each organ forming a common flame. Terminal organs were also stained (albeit weakly and not in all specimens) with antibodies against tyrosinated tubulin (*pnt*, Fig. 2A–C); additionally, tyrosinated tubulin-like immunoreactivity was detected in the protonephridial ducts of some specimens (*pnd*, Fig. 2A–C) revealing that the ducts are anteriorly connected by the loop positioned anteriorly to the brain (*apl*, Fig. 2B).

### Nervous system of the male

Similarly to the females, the nervous system of male *E. senta* consists of a frontal brain, longitudinal nerve cords, coronal and peripheral nerves and sensory structures (Figs. 2D, E, 3C–E, 4D, E; 5C, D). The stomatogastric nervous system is, however, entirely lacking in the males.

The male brain has a similar shape and length to the female’s one (mean length ≈20μm, mean width ≈37μm; N=2), although it is narrower and seems to be more compact (compare Fig. 2 G and H). As in females it is clearly divided into outer layer of perikarya and an internal neuropile (*bp* and *np*, Fig. 2G, respectively). Anteriorly coronal nerves protrude from the brain (*cn*, Figs. 2 D–F, 4D), yet their exact number was difficult to determine due to the aforementioned compactness. Like in females two thick nerves emerge laterally from the brain, one of them continues as an anterior dorsal nerve (*adn*) and connects to the lateral antennae, whereas the other continues as the longitudinal nerve cord (*lnc*) (Figs. 2 D–F, 4D). Dorso-posteriorly two thick nerves (*dan*) connect the neuropile with the dorsal antenna (Figs. 2F, 4D). There are three pairs of SLIR perikarya, which occupy similar positions as those found in the female brain (Figs. 3C–E, 4E), however they are so densely packed that it is impossible to resolve their exact connection with each other. Nevertheless, the anterior SLIR neurites (*an*) and SLIR neurites of longitudinal nerve cords (*lnc*) connect laterally to this cluster of SLIR brain perikarya (Figs. 3C–E, 4E). FMRF-amide-like immunoreactivity was also detected in some of the brain cells (*bp*, Fig. 5C).

Longitudinal nerve cords (*lnc*) extend from the brain to the foot ganglion (Figs. 2D, E; 3C; 4 D, E), but unlike in females they are SLIR along the entire length (Figs. 3C, 4E) but not FLIR. Short foot nerves protrude from the foot ganglion toward the tips of the toes (*fn*, Figs. 4D, 5D). The lateral clusters of weakly FLIR perykarya are present at the anterior portion of the cords (*lp*, Fig. 5C), whereas pair of SLIR perikarya can be detected in the foot ganglion (fg, Figs. 3C, 4E). We did not manage to detect the anterior commissure with any of our immunostainings, but the posterior one exhibits tyrosinated tubulin-like immunoreactivity (*pc*, Figs. 2D, 4D). A pair of fine SLIR neurites extends dorsally from the lateral cords and continues along anterior dorsal nerves, which lead to the lateral antennae (*adn*, Figs. 3C–E, 4E); however, they do not reach the sensory organs themselves. The posterior dorsal nerves were not directly detected, but the supra-anal sensory organs were visible in staining against acetylated and tyrosinated tubulin (*sso*, Figs. 2D, 5D). The weakly tyrosinated tubulin-like immunoreactive (TLIR) nerve innervates each of the supra-anal organs in the male and, even though its connection to the lateral cords was not possible to trace, we assume that it represents the male counterpart of the female posterior dorsal nerve.

Same as in females, five sensory organs are present on the external surface of the male *E. senta*: an unpaired dorsal antenna on the posterior part of the head, the lateral antenna in the middle of the trunk and the supra-anal sensory organs in the posterior part of the trunk (*da, la* and *sso*, Figs. 2D, F; 4D; 5D, respectively). All of those organs seem to have similar arrangement and innervation as their counterparts in female. Four pairs of terminal organs of the male protonephridial system show strong acetylated-TLIR and a typical flame-like organization of cilia (*pnt*, Fig. 5D). Additionally, a strong acetylated-TLIR was detected in the corona, male copulatory organs and in the sperm flagella (*co, mco* and *sp*, Fig. 5D, respectively).

## Discussion

### Differences between females and males of *E. senta*

The nervous systems of both sexes of *E. senta* are similar on both a general and detailed level. The most pronounced difference is related to the complete reduction of stomatogastric nervous system (SNS) in males. Additionally, we find the anterior commissure connecting lateral nerve cords only in females and no corresponding structure was detected in males. Apart from those two structures lacking in males we found counterparts of all female nervous structures in the dwarfed males. Our results are similar to those of the investigation of the sexual dimorphism in musculature of *E. senta* and *Brachionus manjavacas*(11). The males and females have almost identical somatic musculature and differ mostly in the lack of the mastax musculature in males.

The female brain of *E. senta* is approximately two times larger than the male brain by measuring the area of the ellipse appointed by the widest and longest axes of the brain as an indicator of the brain size (area of the widest section of the female brain ≈1093μm^2^, area of the same section in male ≈581μm ^2^, ratio: 1.88). This roughly corresponds with the body size difference between females and males (ratio of the mean body length between sexes: 2.21). However, the cell nuclei in the male brain seem to be more densely packed than in female one, therefore the exact number of cell nuclei in both organs might be actually similar.

Although the general architecture of the nervous system is alike between the two sexes, there are interesting differences in the immunoreactivity of particular structures (Table 1). For instance the lateral nerve cords exhibit FMRF-amide-like immunoreactivity in females but not in males. While on the other hand, their posterior fragments (including the posterior foot ganglion) show serotonin-like immunoreactivity in males but not in females. Further differences in the immunoreactivity are evident for the innervation of lateral antennae, posterior commissure of lateral nerve cords and foot nerves (Table 1). Those differences might indicate that despite similar morphology particular elements of male and female nervous system might vary in their neurophysiology and hence function.

**Table 1.** Summary of the differences between females and males in the immunoreactivity detected in particular morphological structures.

| Structure | Sex | Immunoreactivity |  |  |  |
| --- | --- | --- | --- | --- | --- |
|  |  | Tyrosinated tubulin-like | Acetylated tubulin-like | Serotonin-like | FMRF-amide-like |
| Longitudinal nerve cords | Female | + | - | anterior | + |
|  | Male | + | - | entire | - |
| Posterior commissure | Female | - | - | - | + |
|  | Male | + | - | - | - |
| Anterior dorsal nerve | Female | + | - | - | - |
|  | Male | + | - | + | - |
| Foot ganglion | Female | + | - | - | - |
|  | Male | + | - | + | - |
| Foot nerves | Female | + | - | - | + |
|  | Male | + | + | - | - |

As well as the nervous system we also visualized portion of the excretory organs, including terminal organs (in both sexes) and protonephridial ducts (in females). Both sexes have four pairs of terminal organs with vibratile ciliary flames, which contrasts with two pairs described by Martini based on his LM observation(27). The terminal organs have typical monogonont organization with several cilia forming a common unison flame (e.g.(38)). The movement of those flames was observed in both sexes in LM examination of living specimens. In females we found anterior loop connecting nephridial ducts anteriorly to the brain, the structure known from literature as Huxley’s anastome(1), which has also been described in *E. senta* females(27). The observed details of protonephridia indicate that next to the musculature and nervous system the excretory organs of both sexes are functional and share a similar architecture.

### Male dwarfism in Monogononta

Male dwarfism is a relatively widespread phenomenon present in many organisms(39). Among Spiralia (to which rotifers belong(40-42)) it has been reported in e.g. Cycliophora(43-47), Orthonectida(48), which are now considered parasitic annelids(49, 50), some octopi(51) and in several annelid clades including some dinophilids(52-54), *Osedax*(55-57), Spionidae(58, 59) and bonellid echiurans(60). There are two proposed mechanisms responsible for the origin of dwarfism: progenesis or reduction(61, 62), which are reflected in the morphology of the dwarfed forms, including their nervous system and musculature(52). The progenetic animals resemble earlier (larval or juvenile) developmental stages of normal sized counterparts, whereas dwarfs as a result of reduction lack many characters typical for non-reduced specimens. They have anatomical adaptations to the reduction which do not bear obvious homology to neither larval nor adult structures of the normal-sized specimen(52, 62). Morphology of some of the spiralian dwarfed males resembles the larval arrangement indicating that they emerged through progenesis as in bonellid echiurans and *Osedax*(56, 57, 60), whereas dwarfed males of cycliophorans, orthonectids, Dinophilus gyrociliatus (Dinophilidae) and Scolelepis laonicola (Spionidae) are not similar to the early developmental stages of their female counterparts and rather originated through morphological reduction(45, 48, 52, 59).

At the moment of hatching, the male of *E. senta* is of similar size and complexity as female and only its digestive system with associated structures (mastax musculature, stomatogastric nervous system) is reduced ((11), this study). The post-hatching growth of rotifer females is achieved mostly through increase in the size of the cells but not their number(1, 63), thus the feeding females become larger while their neuroanatomy, musculature and excretory system remains comparable to that of the dwarfed non-feeding male. Hence, with the exception of the digestive tract, the dwarfism of the male results from atrophy, caused by the lack of digestive system, and not from morphological reduction. Instead, the adult male, due to its inability to feed, retains characters of a juvenile hatchling - the situation that can be regarded as a specific case of progenesis.

Both feeding and atrophic males have been reported from monogononts and apparently the species ecology, and not phylogeny, seems to predominantly explain presence of one or the other form(10). This indicates that male atrophy (and subsequent dwarfism) might be reversible in Monogononta. Such evolutionary reversal from dwarfed progenetic male to normal-sized organism was already reported in bone eating annelid *Osedax priapus*, proving that transition to male dwarfism is evolutionarily labile and not necessarily unidirectional(55).

### Nervous system of *E. senta* females – a comparative view

Martini(27) described the morphology of *Epiphanes (=Hydatina)* senta females using LM on intact specimens and histological sections. His description includes, among others, a detailed reconstruction of the nervous system. Results from our investigation show close resemblance with those of Martini. Similarly, Leasi et al.(11) found their CLSM-based reconstruction of musculature congruent with the LM-based reconstruction of Martini. The only neural structure which Martini did not describe and we revealed in our study is the anterior commissure connecting longitudinal nerve cords ventrally to the mastax. This commissure is very thin, does not show tubulin immunoreactivity and was visualized solely with the antibodies against serotonin. Those factors might explain why the structure was omitted in the previous LM-based investigation.

The general neuroarchitecture in monogonont females is quite conserved(1, 20, 26) and our reconstruction of the neuroanatomy of *E. senta* females conforms to the generalized plan of the rotifer nervous system. All of the structures, which we herby described for the female, have been reported in some rotifer species in the previous investigations. There are, however, some aspects of the rotifer nervous system that need an additional discussion.

So far the serotonin and FMRF-amide have been used the most extensively as nervous system markers in rotifers, and comparison of immunoreactivity patterns of those two markers is possible for a broad range of taxa(19-22, 26, 64). Similarly, as in other Monogononta(19, 20, 26), FMRF-amide-like immunoreactivity seems to be more spread than serotonin-like immunoreactivity in the nervous system of *E. senta* females. However, at the same time the exact connectivity of FMRF-amide like immunoreactive perikaryal is impossible to trace, whereas connectivity of serotonin-like perikarya can be reconstructed(20). Therefore, those two markers should be used for different purposes – the first one allows general but imprecise staining of the large portion of the nervous system, whereas the other allows reconstruction of only the small fraction of the system, but with very accurate cellular resolution.

The serotonin-like immunoreactivity in SNS has been reported for all Ploima species investigated thus far(20, 26), but is apparently absent in all examined Gnesiotrocha(19, 21, 22), a discrepancy that has been stressed as an important difference between those two clades(19). However, we did not detect serotonin-like immunoreactivity in the SNS of *E. senta* (which is phylogenetically nested within Ploima), which indicates that serotonin-like immunoreactivity in SNS is a homoplastic character in monogonont rotifers similarly to closely related Gnathostomulida(65).

In the available literature there is also some disagreement regarding connection between SNS and central nervous system in Rotifera. In the older literature the stomatogastric nerve has been described as directly connecting to the brain (e.g. (27, 66)), an arrangement which has been confirmed by Hochberg(20) in his CLSM study on *Notommata copeus*. On the other hand the alternative connection to the lateral nerve cords has been also reported in *N. copeus* and *Asplanchna herricki*(20, 26). In our investigation we found a thick stomatogastric nerve connecting the mastax ganglion directly to the ventro-posterior brain of the female of *E. senta* without evidence of the connection between SNS and longitudinal cords. The pharynx-related ganglion (or at least condensation of neuronal perikarya(67)) connecting directly to the brain has also been reported in other Gnathifera, i.e. Gnathostomulida(65) and Micrognathozoa (where the exact connection of the ganglion to the brain has not been clearly demonstrated(68)) as well as in Chaetognatha(69). According to the recent phylogenies Gnathifera and Chaethognatha seem to form a clade(41), and presence of the pharyngeal ganglion directly connecting to the brain has been already proposed as autapomorphy of Chaetognatha+Gnathifera(65).

## Conclusions

We provide a CLSM-based description of the sex-related differences in the nervous system of the monogonont rotifer, exemplified by the well-studied Epiphanes senta. The neuroanatomy of both sexes is congruent and shows similar level of complexity, though the male nervous system is more compact and lack stomatogastric part due to the reduction of the digestive tract. Additionally some of the nervous structures display different immunoreactivity between sexes possibly indicating divergence in neurophysiology and function. Comparison of nervous system, musculature and excretory organs between feeding females and dwarfed males suggest that male dwarfism originate from progenesis and small body size of males is a result of the growth arrest at the hatchling stage due to the reduction of digestive system and subsequent atrophy.

## Methods

### Animals culturing and fixation

The animals were ordered from a commercial provider of aquatic microinvertebrates (www.sciento.co.uk) in September 2015 and cultured in Jaworski’s medium at 20°C and a 14:10⍰h light:dark cycle. The medium was refreshed every two weeks and the animals were fed ad libitum with the algae *Rhodomonas* sp., *Cryptomonas* sp., and *Chlamydomonas reinhardtii*. Under those conditions both females and males are present in the cultures so there is no need for induction of mixis.

The individual animals were transferred with pipette from cultures to an embryo dish with Jaworski medium; feeding females were starved over night. Prior to fixation, the animals were relaxed for approximately 10 minutes with a solution of 1% bupivacaine and 10% ethanol in culturing medium. Thereafter they were fixed for 1 h in 4% paraformaldehyde solution at room temperature and subsequently rinsed several times with phosphate buffered saline (PBS) with 0.1% Tween-20.

### Immunohistochemistry

After several washes in PBT (PBS + 0.1% Tween-20 + 0.1% bovine serum albumin) animals were preincubated for 30 min at room temperature in PTx+NGS (5% Normal Goat Serum in PBS + 0.1% Triton X-100) and then incubated overnight at 4°C in primary antibodies (mouse anti acetylated tubulin, Sigma T6793 or mouse anti tyrosinated tubulin, Sigma T9028 and rabbit anti serotonin, Sigma S5545 or rabbit anti FMRF-amide, Immunostar 20091) dissolved in PTx+NGS in 1:500 concentrations. The animals were then rinsed several times in PBT, preincubated for 30 min at room temperature in PTX+NGS and incubated overnight at 4°C in secondary antibodies (goat anti-mouse conjugated with AlexaFluor647 and goat anti-rabbit conjugated with AlexaFluor488, Life Technologies) dissolved in Ptx+NGS in 1:250 concentrations. Eventually, the animals were rinsed several times in PBT, stained for cell nuclei with DAPI (1:1000 solution in PBS for 40 minutes) and mounted in 80% glycerol.

Altogether 19 specimens were investigated – six males (three with antibodies against tyrosinated tubulin and serotonin and three with antibodies against acetylated tubulin and FMRF-amide) and 13 females (seven with antibodies against tyrosinated tubulin and serotonin and six with antibodies against acetylated tubulin and FMRF-amide).

### Microscopy and image processing

Mounted specimens were scanned in Leica SP5 confocal laser scanning microscope. Z-stacks of scans were projected into 2D images and 3D reconstructions in IMARIS 9.1.2, which was also used to conduct all the measurements. Schematic drawings based on Z-stacks of scans were made in Adobe Illustrator CS6. Additionally, some living animals anesthetized with bupivacaine solution were photographed with Zeiss Axiocam HRc connected to a Zeiss Axioscope Ax10 using bright-field Nomarski optics. CLSM and light microscopy images were adjusted in Adobe Photoshop CC 2015 and assembled in Adobe Illustrator CS6.

## Declarations

### Ethics approval and consent to participate

Studies of rotifers do not require ethics approval or consent to participate.

### Consent for publication

Not applicable.

### Availability of data and material

All data generated or analyzed during this study are included in this published article.

## Authors’ contributions

AF and LG kept animal cultures, fixed animals and performed antibody staining and confocal imaging. AH designed the study and contributed to writing. LG arranged figures and drafted manuscript. All authors read and accepted the final version of the manuscript.

## Acknowledgements

We thank the team of the Sars Group “Comparative Developmental Biology” for help and discussions and particularly Daniel Thiel for his help with RF-amide staining.

## Additional file 1

COX1 sequences retrived from the transcriptome of investigated *Epiphanes* cf. *senta* in FASTA format:

>E._cf._senta_COX1_seq1

CTTCTACAAATCATAAGGATATTGGTACTCTTTATTTTATCTTTGGTATGTGAGCTGGGT TTATTGGTCTTAGAATGAGTTTACTTATTCGTTTAGAGCTTGGGGTTGTAGGCCCCTACC TTGGTGATGAGCATCTTTATAATGTTTTAGTTACGGCTCATGCTTTTGTTATGATTTTCT TTATAGTTATACCTGTGTCTATGGGTGGTTTTGGTAATTGACTAATTCCTCTTATGTTAG GTGTTGCTGATATAGCTTTTCCTCGTATAAACAATTTATCTTTCTGACTTTTAGTTCCTT CTTTTCTTTTCCTACTTCTATCTTCTATTTTAGATGCGGGAGCAGGTACTGGGTGAACTG TTTATCCTCCTCTTTCTGATTCTAAGTATCATAGTGGTATTTCTGTAGATCTTGCTATTT TTAGGTTACATCTAGCTGGTGTTTCTTCTATTTTAGGTAGAATTAATTTCCTTACAACTA TTATTTGTTCTCGTACTACTAAAGCAGTTTCTCTTGATCGTCTTCCTCTAATGCTTTGGG CTATTGCAGTTACCGCAGTTTTATTAGTTACTAGACTTCCTGTTCTTGCAGGAGCTATCA CTATGCTTTTAACTGATCGTAATTTTAACACTTCTTTCTTTGATCCTGCTGGTGGTGGTA ACCCTGTTTTATATCAACATCTTTTT

>E._cf._senta_COX1_seq2

CTTCTACAAATCATAAGGATATTGGTACTCTTTATTTTATCTTTGGTATGTGAGCTGGGT TTATTGGTCTTAGAATGAGTTTACTTATTCGTTTAGAGCTTGGGGTTGTAGGCCCCTACC TTGGTGATGAGCATCTTTATAATGTTTTAGTTACGGCTCATGCTTTTGTTATGATTTTCT TTATAGTTATACCTGTGTCTATGGGTGGTTTTGGTAATTGACTAATTCCTCTTATGTTAG GTGTTGCTGATATAGCTTTTCCTCGTATAAACAATTTATCTTTCTGACTTTTAGTTCCTT CTTTTCTTTTCCTACTTCTATCTTCTATTTTAGATGCGGGAGCAGGTACTGGGTGAACTG TTTATCCTCCTCTTTCTGATTCTAAGTATCATAGTGGTATTTCTGTAGATCTTGCTATTT TTAGGTTACATCTAGCTGGTGTTTCTTCTATTCTAGGCAGAATTAATTTCCTTACAACTA TTATTTGTTCTCGTACTACTAAAGCAGTTTCTCTTGATCGTCTTCCTCTTATGCTTTGGG CTATTGCAGTTACTGCAGTTTTATTAGTTACTAGACTTCCTGTTCTTGCAGGAGCTATCA CTATGCTTTTAACTGATCGTAATTTTAACACTTCTTTCTTTGATCCTGCTGGTGGTGGTA ATCCTGTTCTATATCAACATCTTTTT

## References

1. Fontaneto D, De Smet W. 4. Rotifera. Handbook of zoology, Gastrotricha and Gnathifera. 2015:217–300.

2. Serra M, Snell TW. Sex loss in monogonont rotifers. Lost Sex: Springer; 2009. p. 281–94.

3. Leasi F, Ricci C. Musculature of two bdelloid rotifers, Adineta ricciae and Macrotrachela quadricornifera: organization in a functional and evolutionary perspective. J Zool Syst Evol Res. 2010;48(1):33–9.

4. Serra M, Snell TW. Why are male rotifers dwarf? Trends Ecol Evol. 1998;13(9):360–1.

5. Birky Jr CW, Gilbert JJ. Parthenogenesis in rotifers: the control of sexual and asexual reproduction. American Zoologist. 1971;11(2):245–66.

6. Pourriot R, Clément P. Action de facteurs externes sur la reproduction et le cycle reproducteur des Rotifers. Acta Oecologica Generale. 1981;2:135–51.

7. Snell TW. A review of the molecular mechanisms of monogonont rotifer reproduction. Hydrobiologia. 2011;662(1):89–97.

8. Stelzer CP, Snell TW. Induction of sexual reproduction in Brachionus plicatilis (Monogononta, Rotifera) by a density-dependent chemical cue. Limnol Oceanogr. 2003;48(2):939–43.

9. Schröder T, Walsh EJ. Genetic differentiation, behavioural reproductive isolation and mixis cues in three sibling species of Monogonont rotifers. Freshwater Biol. 2010;55(12):2570–84.

10. Ricci C, Melone G. Dwarf males in monogonont rotifers. Aquatic Ecology. 1998;32(4):361–5.

11. Leasi F, Fontaneto D, Melone G. Phylogenetic constraints in the muscular system of rotifer males: investigation on the musculature of males versus females of Brachionus manjavacas and Epiphanes senta (Rotifera, Monogononta). J Zool. 2010;282(2):109–19.

12. Keshmirian J, Nogrady T. Histofluorescent Labeling of Catecholaminergic Structures in Rotifers (Aschelminthes). 2. Males of Brachionus plicatilis and Structures from Sectioned Females. Histochemistry. 1988;89(2):189–92.

13. Clément PW, E. Rotifera. In:Harrison FWR, E. E., editor. Microscopic anatomy of invertebrates. 4: Aschelminthes. New York: Wiley-Liss; 1991. p. 219–97.

14. Dehl E. Morphologie von Lindia tecusa Zeitschrift für wissenschaftliche Zoologie 1934;145:169–219.

15. Eakin RM, Westfall JA. Ultrastructure of the Eye of the Rotifer Asplanchna brightwelli. J Ultrastruct Res. 1965;12:46–62.

16. Hirschfelder G. Beiträge zur Histologie der Rädertiere (Eosphora, Hydatina, Euchlanis, Notommata). Zeitschrift für wissenschaftliche Zoologie. 1910;96:209–335.

17. Hlava S. Beiträge zur Kenntnis der Rädertiere: über die Anatomie von Conochiloides natans. Zeitschrift für wissenschaftliche Zoologie. 1905;80:282–326.

18. Hochberg R. Three-dimensional reconstruction and neural map of the serotonergic brain of Asplanchna brightwellii (Rotifera, Monogononta). Journal of morphology. 2009;270(4):430–41.

19. Hochberg R. On the serotonergic nervous system of two planktonic rotifers, Conochilus coenobasis and C. dossuarius (Monogononta, Flosculariacea, Conochilidae). Zoologischer Anzeiger. 2006;245(1):53–62.

20. Hochberg R. Topology of the nervous system of Notommata copeus (Rotifera: Monogononta) revealed with anti-FMRFamide, -SCPb, and -serotonin (5-HT) immunohistochemistry. Invertebrate Biology. 2007;126(3):247–56.

21. Hochberg R, Lilley G. Neuromuscular organization of the freshwater colonial rotifer, Sinantherina socialis, and its implications for understanding the evolution of coloniality in Rotifera. Zoomorphology (Berlin). 2010;129(3):153–62.

22. Hochberg A, Hochberg R. Serotonin immunoreactivity in the nervous system of the free-swimming larvae and sessile adult females of Stephanoceros fimbriatus (Rotifera: Gnesiotrocha). Invertebrate Biology. 2015;134(4):261–70.

23. Keshmirian J, Nogrady T. Histofluorescent labelling of catecholaminergic structures in rotifers (Aschelminthes) in whole animals. Histochemistry. 1987;87(4):351–7.

24. Kotikova EA. Localization and neuroanatomy of catecholaminergic neurons in some rotifer species. Hydrobiologia. 1995;313-314:123–7.

25. Kotikova EA. Catecholaminergic neurons in the brain of rotifers. Hydrobiologia. 1998;387-388:135–40.

26. Kotikova EA, Raikova OI, Reuter M, Gustafsson MKS. Rotifer nervous system visualized by FMRFamide and 5-HT immunocytochemistry and confocal laser scanning microscopy. Hydrobiologia. 2005;546:239–48.

27. Martini E. Studien über die Konstanz histologischer Elemente. III. Hydatina senta. Zeitschrift für wissenschaftliche Zoologie. 1912;102(425):645.

28. Nachtwey R. Untersuchungen über die Keimbahn, Organogenese und Anatomie von Asplanchna priodonta Gosse. Zeitschrift für wissenschaftliche Zoologie. 1925;126:239–492.

29. Nogrady T, Alai M. Cholinergic neurotransmission in rotifers.Hydrobiologia. 1983;104:149–53.

30. Peters W. Untersuchungen über Anatomie und Zellkonstanz von Synchaeta (S. grimpei Rem., S. baltica Ehrb., S. tavina Hood und S. triophthalma Laut.). Zeitschrift für wissenschaftliche Zoologie. 1931;139:1–119.

31. Remane A. Aschelminthes. Rotatoria. Bronn’s Klassen und Ordnungen des Tier-Reichs. 4. Leipzig: Akademische Verlagsgesellschaft; 1933. p. 1–577.

32. Seehaus W. Zur Morphologie der Rädertiergattung Testudinella Bory de St. Vincent (= Pterodina Ehrenberg). Zeitschrift für wissenschaftliche Zoologie. 1930;137:175–273.

33. Stossberg K. Zur Morphologie der Rädertiergattungen Euchlanis, Brachionus und Rhinoglena. Zeitschrift für wissenschaftliche Zoologie. 1932;142:313–424.

34. Martin-Duran JM, Pang K, Børve A, Le HS, Furu A, Cannon JT, et al. Convergent evolution of bilaterian nerve cords. Nature. 2018;553(7686):45-+.

35. Fröbius AC, Funch P. Rotiferan Hox genes give new insights into the evolution of metazoan bodyplans. Nat Commun. 2017;8.

36. Schröder T, Walsh EJ. Cryptic speciation in the cosmopolitan Epiphanes senta complex (Monogononta, Rotifera) with the description of new species. Hydrobiologia. 2007;593:129–40.

37. Schröder T. Precopulatory mate guarding and mating behaviour in the rotifer Epiphanes senta (Monogononta: Rotifera). P Roy Soc B-Biol Sci. 2003;270(1527):1965–70.

38. Riemann O, Ahlrichs WH. The evolution of the protonephridial terminal organ across Rotifera with particular emphasis on Dicranophorus forcipatus, Encentrum mucronatum and Erignatha clastopis (Rotifera: Dicranophoridae). Acta Zoologica (Copenhagen). 2010;91(2):199–211.

39. Vollrath F. Dwarf males. Trends Ecol Evol. 1998;13(4):159–63.

40. Laumer CE, Bekkouche N, Kerbl A, Goetz F, Neves RC, Sørensen MV, et al. Spiralian phylogeny informs the evolution of microscopic lineages. Current Biology. 2015;25(15):2000–6.

41. Marlétaz F, Peijnenburg KTCA, Goto T, Satoh N, Rokhsar DS. A New Spiralian Phylogeny Places the Enigmatic Arrow Worms among Gnathiferans. Current Biology. 2019;29(2):312-+.

42. Hejnol A. A twist in time—the evolution of spiral cleavage in the light of animal phylogeny. Integrative and comparative biology. 2010;50(5):695–706.

43. Neves RC, da Cunha MR, Funch P, Wanninger A, Kristensen RM. External morphology of the cycliophoran dwarf male: a comparative study of Symbion pandora and S. americanus. Helgoland Marine Research. 2010;64(3):257–62.

44. Neves RC, Kristensen RM, Wanninger A. Serotonin immunoreactivity in the nervous system of the Pandora larva, the Prometheus larva, and the dwarf male of Symbion americanus (Cycliophora). Zoologischer Anzeiger. 2010;249(1):1–12.

45. Neves RC, Reichert H. Microanatomy and Development of the Dwarf Male of Symbion pandora (Phylum Cycliophora): New Insights from Ultrastructural Investigation Based on Serial Section Electron Microscopy. PLoS ONE. 2015;10(4):e0122364.

46. Neves RC, Sørensen KJK, Kristensen RM, Wanninger A. Cycliophoran Dwarf Males Break the Rule: High Complexity with Low Cell Numbers. Biol Bull-Us. 2009;217(1):2–5.

47. Obst M, Funch P. Dwarf male of Symbion pandora (Cycliophora). Journal of Morphology. 2003;255(3):261–78.

48. Slyusarev GS, Nesterenko MA, Starunov VV. The structure of the muscular and nervous systems of the male Intoshia linei (Orthonectida). Acta Zool-Stockholm.0(0).

49. Bondarenko N, Bondarenko A, Starunov V, Slyusarev G. Comparative analysis of the mitochondrial genomes of Orthonectida: insights into the evolution of an invertebrate parasite species. Molecular Genetics and Genomics. 2019.

50. Schiffer PH, Robertson HE, Telford MJ. Orthonectids Are Highly Degenerate Annelid Worms. Current Biology. 2018;28(12):1970-4.e3.

51. Norman MD, Paul D, Finn J, Tregenza T. First encounter with a live male blanket octopus: The world’s most sexually size-dimorphic large animal. New Zealand Journal of Marine and Freshwater Research. 2002;36(4):733–6.

52. Kerbl A, Fofanova EG, Mayorova TD, Voronezhskaya EE, Worsaae K. Comparison of neuromuscular development in two dinophilid species (Annelida) suggests progenetic origin of Dinophilus gyrociliatus. Front Zool. 2016;13.

53. Windoffer R, Westheide W. The Nervous system of the male Dinophilus gyrociliatus (Polychaeta, Dinophilidae).2. Electron microscopical reconstruction of nervous anatomy and effector cells. J Comp Neurol. 1988;272(4):475–88.

54. Windoffer R, Westheide W. The nervous system of the male Dinophilus gyrociliatus (Annelida, Polychaeta).1. Number, types and distribution pattern of sensory cells. Acta Zool-Stockholm. 1988;69(1):55–64.

55. Rouse GW, Wilson NG, Worsaae K, Vrijenhoek RC. A dwarf male reversal in bone-eating worms. Curr Biol. 2015;25(2):236–41.

56. Rouse GW, Worsaae K, Johnson SB, Jones WJ, Vrijenhoek RC. Acquisition of dwarf male “harems” by recently settled females of Osedax roseus n. sp. (Siboglinidae; Annelida). Biol Bull. 2008;214(1):67–82.

57. Worsaae K, Rouse GW. The simplicity of males: dwarf males of four species of Osedax (Siboglinidae; Annelida) investigated by confocal laser scanning microscopy. J Morphol. 2010;271(2):127–42.

58. Vortsepneva E, Tzetlin A, Purschke G, Mugue N, Hass-Cordes E, Zhadan A. The parasitic polychaete known as Asetocalamyzas laonicola (Calamyzidae) is in fact the dwarf male of the spionid Scolelepis laonicola (comb. nov.). Invertebrate Biology. 2008;127(4):403–16.

59. Vortsepneva E, Tzetlin A, Tsitrin E. Nervous system of the dwarf ectoparasitic male of Scolelepis laonicola (Polychaeta, Spionidae). Zoosymposia. 2009;2:437–45.

60. Schuchert P, Rieger RM. Ultrastructural observations on the dwarf male of Bonellia viridis (Echiura). Acta Zoologica (Copenhagen). 1990;71(1):5–16.

61. Gould SJ. Ontogeny and Phylogeny. Cambridge, MA: Bel-knap Press of Harvard Universitiy Press; 1977.

62. Hanken J, Wake DB. Miniaturization of body size: organismal consequences and evolutionary significance. Annual Review of Ecology and Systematics. 1993;24:501–19.

63. Hejnol A. Gnathifera. Evolutionary Developmental Biology of Invertebrates 2: Springer; 2015. p. 1–12.

64. Leasi F, Pennati R, Ricci C. First description of the serotonergic nervous system in a bdelloid rotifer: Macrotrachela quadricornifera Milne 1886 (Philodinidae). Zoologischer Anzeiger. 2009;248(1):47–55.

65. Gąsiorowski L, Bekkouche N, Worsaae K. Morphology and evolution of the nervous system in Gnathostomulida (Gnathifera, Spiralia). Organisms Diversity & Evolution. 2017;17(1):447–75.

66. Hyman LH. IV. Class Rotifera. In: Hyman LH, editor. The invertebrates: Acanthocephala, Aschelminthes, and Entoprocta The pseudocoelomate Bilateria. Vol. III. New Yor, Toronto, London: McGraw-Hill Book Company, Inc.; 1951. p. 59–151.

67. Herlyn H. Enigmatic Gnathostomulida (Gnathifera, Spiralia): about monociliated pharyngeal receptors and the pharyngeal nervous system. Zoomorphology (Berlin). 2017;136(4):425–34.

68. Bekkouche N, Worsaae K. Nervous system and ciliary structures of Micrognathozoa (Gnathifera): evolutionary insight from an early branch in Spiralia. Royal Society Open Science. 2016;3(10):160289.

69. Rieger V, Perez Y, Mueller CHG, Lipke E, Sombke A, Hansson BS, et al. Immunohistochemical analysis and 3D reconstruction of the cephalic nervous system in Chaetognatha: insights into the evolution of an early bilaterian brain? Invertebrate Biology. 2010;129(1):77–104.

